# PIEZO1-mediated mechanosensing governs NK cell killing efficiency and infiltration in three-dimensional matrices

**DOI:** 10.1101/2023.03.27.534435

**Authors:** Archana K. Yanamandra, Jingnan Zhang, Galia Montalvo, Xiangda Zhou, Doreen Biedenweg, Renping Zhao, Shulagna Sharma, Markus Hoth, Franziska Lautenschläger, Oliver Otto, Aránzazu del Campo, Bin Qu

**Affiliations:** Biophysics, Center for Integrative Physiology and Molecular Medicine (CIPMM), School of Medicine, Saarland University, Homburg, Germany; INM - Leibniz Institute for New Materials, Saarbrücken, Germany; Department of Experimental Physics, Saarland University, Saarbrücken, Germany; Center for Biophysics, Saarland University, Saarbrücken, Germany; Institute of Physics, University of Greifswald, Greifswald, Germany; Chemistry Department, Saarland University, Saarbrücken, Germany

**Keywords:** Natural killer cells, mechanosensing, PIEZO1, killing efficiency, 3D matrices

## Abstract

Natural killer (NK) cells play a vital role in eliminating tumorigenic cells. Efficient locating and killing of target cells in complex three-dimensional (3D) environments are critical for their functions under physiological conditions. However, the role of mechanosensing in regulating NK cell killing efficiency in physiologically relevant scenarios is poorly understood. Here, we report that the responsiveness of NK cells is regulated by tumor cell stiffness. NK cell killing efficiency in 3D is impaired against softened tumor cells, while it is enhanced against stiffened tumor cells. Notably, the durations required for NK cell killing and detachment are significantly shortened for stiffened tumor cells. Furthermore, we have identified PIEZO1 as the predominantly expressed mechanosensitive ion channel among the examined candidates in NK cells. Perturbation of PIEZO1 abolishes stiffness-dependent NK cell responsiveness, significantly impairs the killing efficiency of NK cells in 3D, and substantially reduces NK cell infiltration into 3D collagen matrices. Conversely, PIEZO1 activation enhances NK killing efficiency as well as infiltration. In conclusion, our findings demonstrate that PIEZO1-mediated mechanosensing is crucial for NK killing functions, highlighting the role of mechanosensing in NK cell killing efficiency under 3D physiological conditions and the influence of environmental physical cues on NK cell functions.

## Introduction

Natural killer (NK) cells belong to the innate immune system, responsible for eliminating aberrant cells such as tumorigenic cells and pathogen-infected cells. In both physiological and pathological conditions, NK cells must navigate through three-dimensional (3D) environments to locate their target cells. NK cells identify the cognate target cells through the engagement of their activating receptors with ligands on the target cell surface and/or detection of absence of self-molecules using their inhibitory receptors (1). Upon target cell recognition, NK cells form an intimate contact termed immunological synapse (IS) and reorient the killing machineries towards target cells (2). The primary killing mechanism employed by NK cells is lytic granules containing cytotoxic proteins such as pore-forming protein perforin and serine protease granzymes. Lytic granules are enriched and released at the IS to induce apoptosis or direct lysis of target cells (3). The release of lytic granules, also known as degranulation, is a hall mark of NK activation triggered by target cell recognition.

Stiffness is a physical characteristic that can differ significantly between healthy and diseased tissues, and stiffness at the tissue level and cell level can differ significantly. For instance, solid tumors are often stiffer than the neighboring healthy tissues primarily due to a highly compacted extracellular matrix (4). Conversely, malignant cells with a high potential for metastasis are typically softer than their counterparts (5, 6). Despite the extensive research on the functional role of chemical cues, the impact of stiffness on functions of immune killer cells, especially in killing-related processes, has only recently gained attention. For NK cells, stiffer substrates potentiate polarization of MTOC, enrichment and release of lytic granules, cytokine production, and the stability of the IS (7). In addition, the actin retrograde flow at the IS, which regulates the NK cell response, is influenced by substrate stiffness (8). The stiffness of cancer cells typically ranges from a few hundred to a few thousand Pa (9-13). The levels of substrate stiffness used to investigate the stiffness-regulated NK cell function is often two to three orders of magnitude higher than the actual stiffness of cancer cells. Therefore, the precise effect of the physiological range of tumor cell stiffness on the effector functions and the corresponding killing efficiency of NK cells remains unclear.

To detect environmental stiffness, cells rely on mechanosensing through surface mechanosensors, mainly mechanically activated ion channels (14). In this regard, the PIEZO family members are the most extensively studied mechanosensors. In T cells, PIEZO1-mediated mechanosensing of fluid shear stress has been found to potentiate T cell activation (15). Additionally, PIEZO1 activation at the IS is essential for optimal T cell receptor signal transduction, potentially through PIEZO1-mediated Ca^2+^ influx (16). In mice, genetic deletion of PIEZO1 in T cells selectively expand Treg population and attenuates the severity of experimental autoimmune encephalomyelitis, an animal model for multiple sclerosis (17). In myeloid cells, PIEZO1-mediated mechanosensing of cyclical pressure, as experienced in lungs, plays a key role in the initiation of proinflammatory response elicited by macrophages and monocytes (18). However, the functional roles of PIEZOs in NK cells have not been characterized.

In this study, we show that the efficiency of NK cell-mediated target cell elimination is regulated by the stiffness of target cells. Specifically, the cytotoxicity of NK cells is decreased against softer target cells and elevated against stiffer target cells. In human NK cells, mechanosensing is primarily mediated by PIEZO1, and perturbation of PIEZO1 abolishes stiffness-dependent responsiveness of NK cells. Furthermore, PIEZO1-mediated mechanosensing governs the infiltration of NK cells into 3D collagen matrices, significantly impacting NK cell killing efficiency in 3D scenarios. In summary, our results highlight the critical regulatory roles of mechanosensing in NK cell-mediated target cell elimination in physiologically relevant 3D scenarios.

## Results

### NK cell activation is regulated by surface stiffness

To investigate how the stiffness of target cells affects NK cell responsiveness, we used functionalized hydrogels of various stiffness as a model system to mimic target cells as reported previously (19). Specifically, we employed poly (acrylamide-co-acrylic acid) (PAAm-co-AA) hydrogels with Young’s Modulus of 2, 12, and 50 kPa functionalized with an activating antibody targeting NKp46 (Fig. 1A), which belongs to the natural cytotoxicity receptor family. The coating efficiency for the hydrogels with these three stiffness levels has been demonstrated the same in our previous work (19). We used NK cells that were isolated from healthy donors and stimulated with IL-2 for 3 days. To evaluate NK activation, we settled NK cells on the functionalized hydrogels at 37°C for four hours and assessed degranulation of lytic granules based on the levels of CD107a on the surface of the NK cells. CD107a is exclusively expressed on vesicular membrane of lytic granules and can only be integrated into the plasma membrane of NK cells after lytic granule release (20). Based on activation-triggered degranulation, we found that NK cells fell into four categories: only activated on 50 kPa (5 out of 12 donors, Fig. 1B), only activated on 12 kPa (3 out of 12 donors, Fig. 1C), activated on all three stiffness levels (2 out of 12 donors, Fig. 1D), or no response (2 out of 12 donors, Fig. 1E). Importantly, degranulation was only triggered by activation of NKp46, as isotype IgG-coated hydrogels did not induce degranulation (Fig. 1B-D, isotype). Notably, NK cells from most donors (10 out of 12) did not respond to very soft hydrogels (2 kPa) (Fig. 1F, G). These findings show that NK cells cannot be fully activated on soft substrates, which is consistent with the reports from the others (7, 8). Based on these results, we hypothesized that softening target cells would impair NK killing capacity.

**Figure 1.**
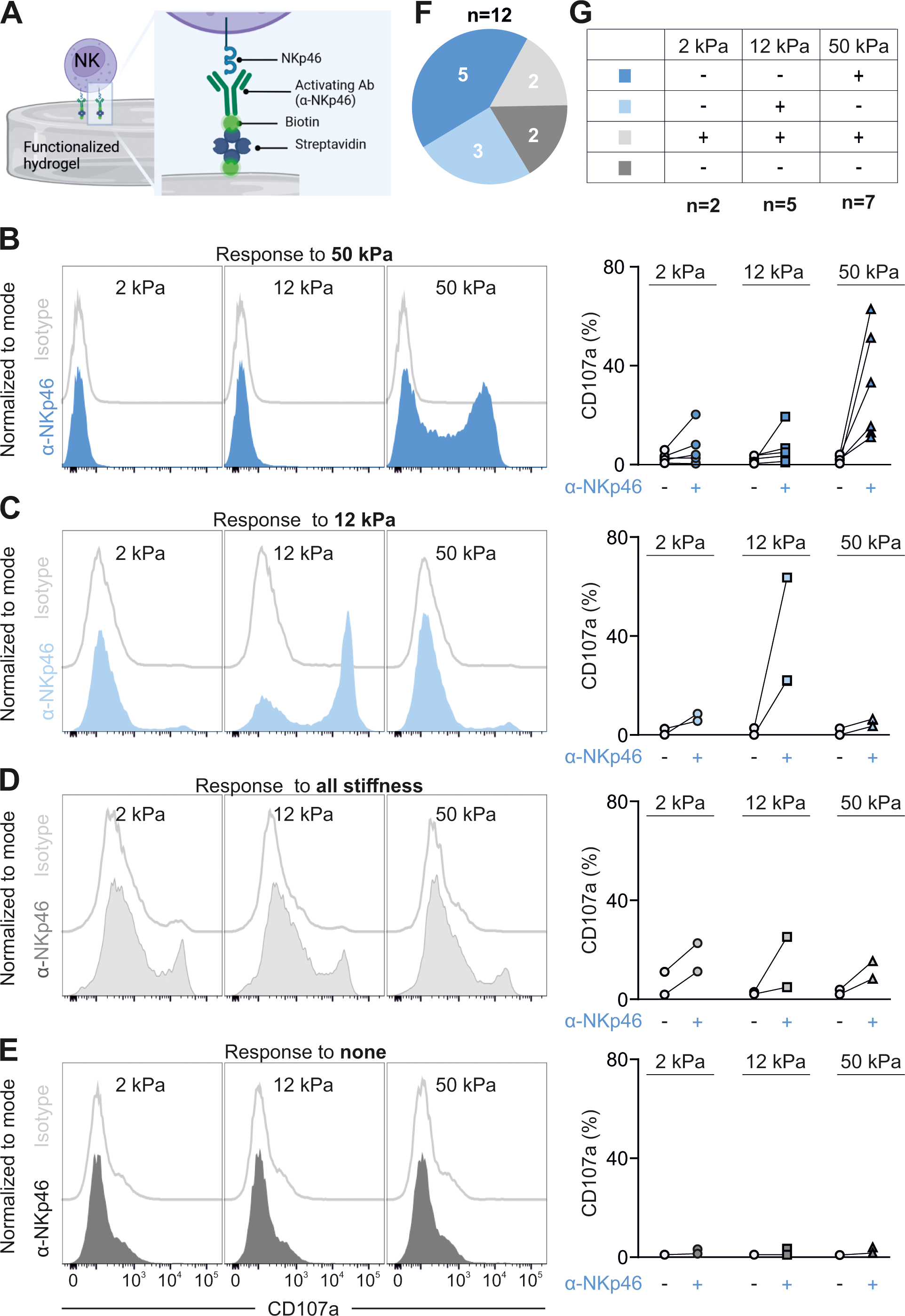
NK cell responsiveness to varying levels of substrate stiffness. Primary human NK cells from healthy donors were stimulated with IL-2 for three days prior to the experiments. (**A**) Sketch of functionalization of hydrogels. PAAm-co-AA hydrogels were first treated with streptavidin and then incubated with biotinylated anti-NKp46 antibodies. (**B-E**) NK cell responsiveness is substrate stiffness-dependent. The activation of NK cells was evaluated using the CD107a degranulation assay. The NK cells were settled on functionalized hydrogels at 37°C with 5% CO_2_ for 4 hours in the presence of anti-CD107a antibody and Golgi Stop. The samples were then analyzed using flow cytometry. The NK cells responded to 50 kPa (**B**, n=5), 12 kPa (**C**, n=3), all stiffness levels (**D**, n=2), or did not respond to any stiffness (**E**, n=2). One representative donor is shown in the left panel and the quantification of all donors is shown in the right panel. (**F-G**) Summary of the NK cell responsiveness from different donors.

### Target cell stiffness modulates NK cell cytotoxicity

To test this hypothesis, we softened the target cells (K562 cells) by DMSO treatment as determined by real-time deformability cytometry (RT-DC) (Fig. 2A). The K562 cells used in our study stably express a FRET-based apoptosis reporter pCasper (K562-pCasper), consisting of a GFP and RFP pair linked by a caspase recognition site (DEVD) (21). Upon initiation of apoptosis, the orange-colored target cells lose their FRET signal and turn green. In the case of necrosis, fluorescent proteins would leak out of the destructed plasma membrane, resulting in the complete loss of fluorescence. To evaluate NK cell killing efficiency in a 3D environment, we embedded K562-pCasper target cells in bovine type I collagen and added IL-2-stimulated primary human NK cells from the top after solidification. This setup allows NK cells to infiltrate the collagen matrix and search for their target cells in a physiologically relevant scenario. We monitored killing events at 37°C every 20 minutes for 48 hours using a high content imaging system. Our results show that the killing efficiency of NK cells against softened target cells was reduced compared to that against the control group (Fig. 2B, Supplementary Movie 1). These findings suggest that softening of tumor cells weakens NK cell killing capacity.

**Figure 2.**
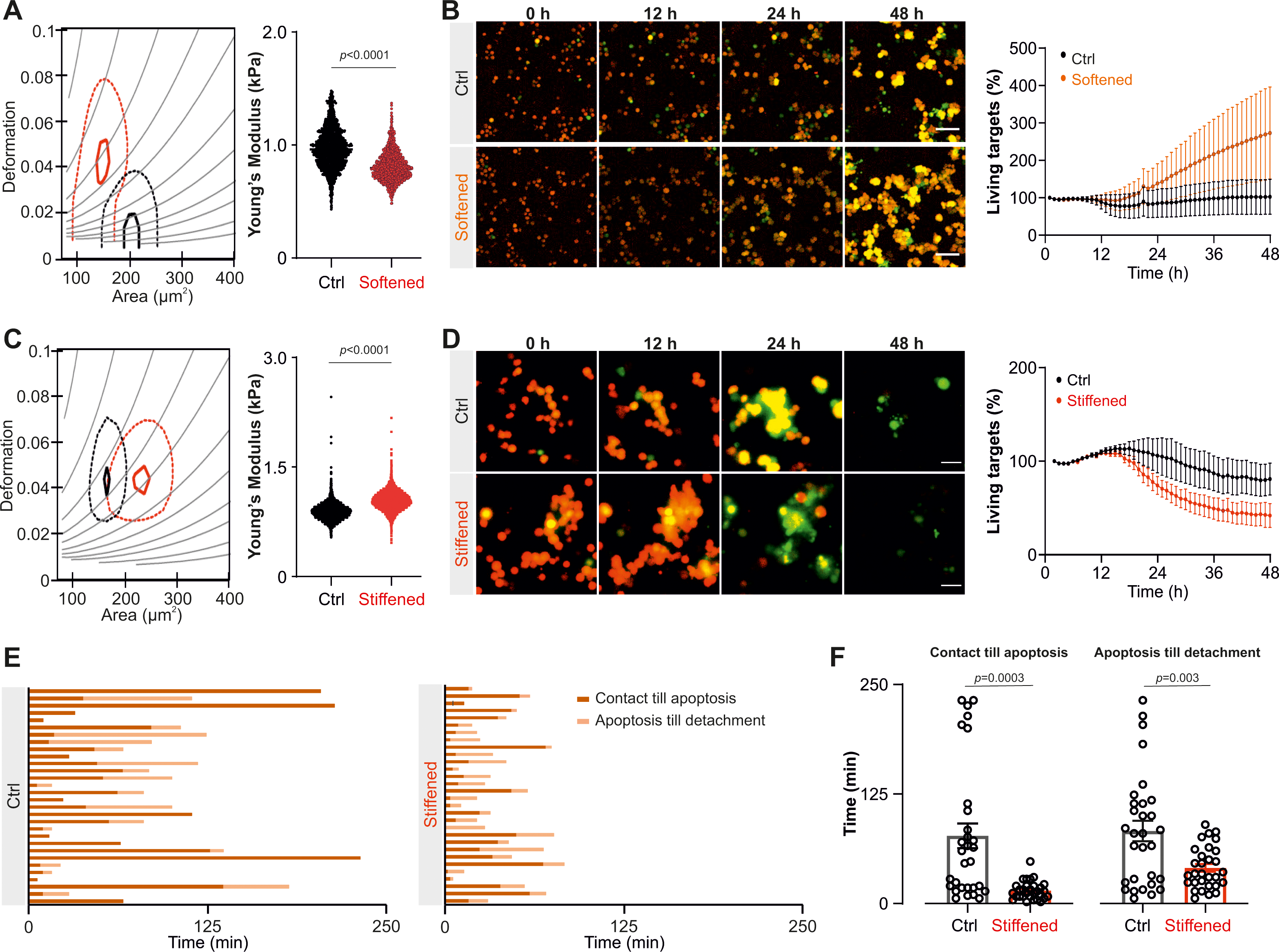
The killing efficiency of NK cells in 3D scenarios is regulated by tumor cell stiffness. Primary human NK cells from healthy donors were stimulated with IL-2 for three days prior to the experiments. K562-pCasper target cells were embedded in collagen matrices (2 mg/ml) and the NK cells were added from the top. Live target cells are in orange-yellow color and apoptotic target cells in green. (**A-B**) Softening tumor cells impairs NK cell killing efficiency in 3D. K562-pCasper cells were pre-treated with DMSO (1:2000, Softened) for 12 hours. Their stiffness was determined using RT-DC (**A**). Time lapse of killing events obtained in 10x magnification and the quantification is shown in **B**. (**C-D**) NK cells eliminate stiffened tumor cells more efficiently. K562-pCasper cells were pre-treated with Blebbistatin (50 µM, Stiffened) for 12 hours. Their stiffness was determined using RT-DC (**C**). Statistical analysis for RT-DC was done using linear mixed models. Time lapse of killing events obtained in 20x magnification and the quantification is shown in **D**. (**E-F**) The duration required for NK cell killing and detachment from the stiffened tumor cells is shortened. K562-pCasper target cells were treated with DMSO (Ctrl) or Blebbistatin (Stiffened). The NK cells were co-incubated with target cells for 4 hours. The NK cells were tracked manually. The duration required for each killing event (the time from the initiation of NK/target contact to target cell apoptosis) and the duration required for NK cell detachment (the time from the initiation of NK/target contact to disengagement of NK cells from the targets) for all NK cells analyzed are shown in the left and right panel of **E**, respectively. The quantification of these durations is shown in **F**. For statistical analysis, the Mann-Whitney-U-test was used. The results were from at least three independent experiments. The data are presented as mean ± SEM. Scale bars are 40 µm.

Next, we examined whether increasing the stiffness of tumor cells could have the opposite effect. To do this, we used blebbistatin, a myosin IIA inhibitor known to enhance the stiffness of cells in suspension by perturbing actomyosin contractility (22). Our analysis of RT-DC revealed that blebbistatin treatment reduced the deformability of target cells, indicating an increase in their stiffness relative to vehicle-treated control cells (Fig. 2C). Consistent with the postulation, we observed an elevated killing efficiency of NK cells against the stiffened blebbistatin-treated tumor cells (Fig. 2D, Supplementary Movie 2). Notably, treatment with DMSO or blebbistatin exhibited no impact on proliferation kinetics of the K562-pCasper cells (Supplementary Fig. 1) and the altered stiffness could persist (Supplementary Fig. 2), reinforcing our conclusion that the observed changes in killing efficiency against DMSO- or Blebbistatin-treated target cells are not a result of alterations in proliferation kinetics per se, but rather attributable to changes in target cell stiffness. Furthermore, analysis of live cell imaging showed a significant reduction in the time required for NK killing (i.e., duration from contact to apoptosis) and the total contact time between NK and target cells (i.e., duration from contact till detachment) in the case of stiffened tumor cells compared to the control group (Fig. 2E, F). Together, these results suggest that the stiffness of target cells has a significant impact on the outcome of NK killing efficiency.

### PIEZO1 mediates NK cell responsiveness to target cell stiffness

Mechano-sensing is crucial for cells to detect stiffness of surrounding environment and the cells they encounter. Among the mechanosensitive channels, PIEZO1 is the most predominantly expressed in primary human NK cells (Fig. 3A). Both unstimulated and stimulated NK cells expressed high levels of PIEZO1 protein, with the majority (>95%) of NK cells expressing PIEZO1 (Supplementary Fig. 3A). PIEZO1 is present on the plasma membrane, exhibiting a distribution pattern similar to F-actin (Supplementary Fig. 3B). To examine the functional role of PIEZO1 in stiffness-regulated NK activation, we used GsMTx4, a peptide isolated from spider venom that inhibits the mechanosensitivity of PIEZO1 (23). Our results show that the surface stiffness-dependent degranulation of NK cells was completely abolished by GsMTx4 treatment (Fig. 3B), indicating that PIEZO1 plays a pivotal role in mediating mechano-sensing in NK cells.

**Figure 3.**
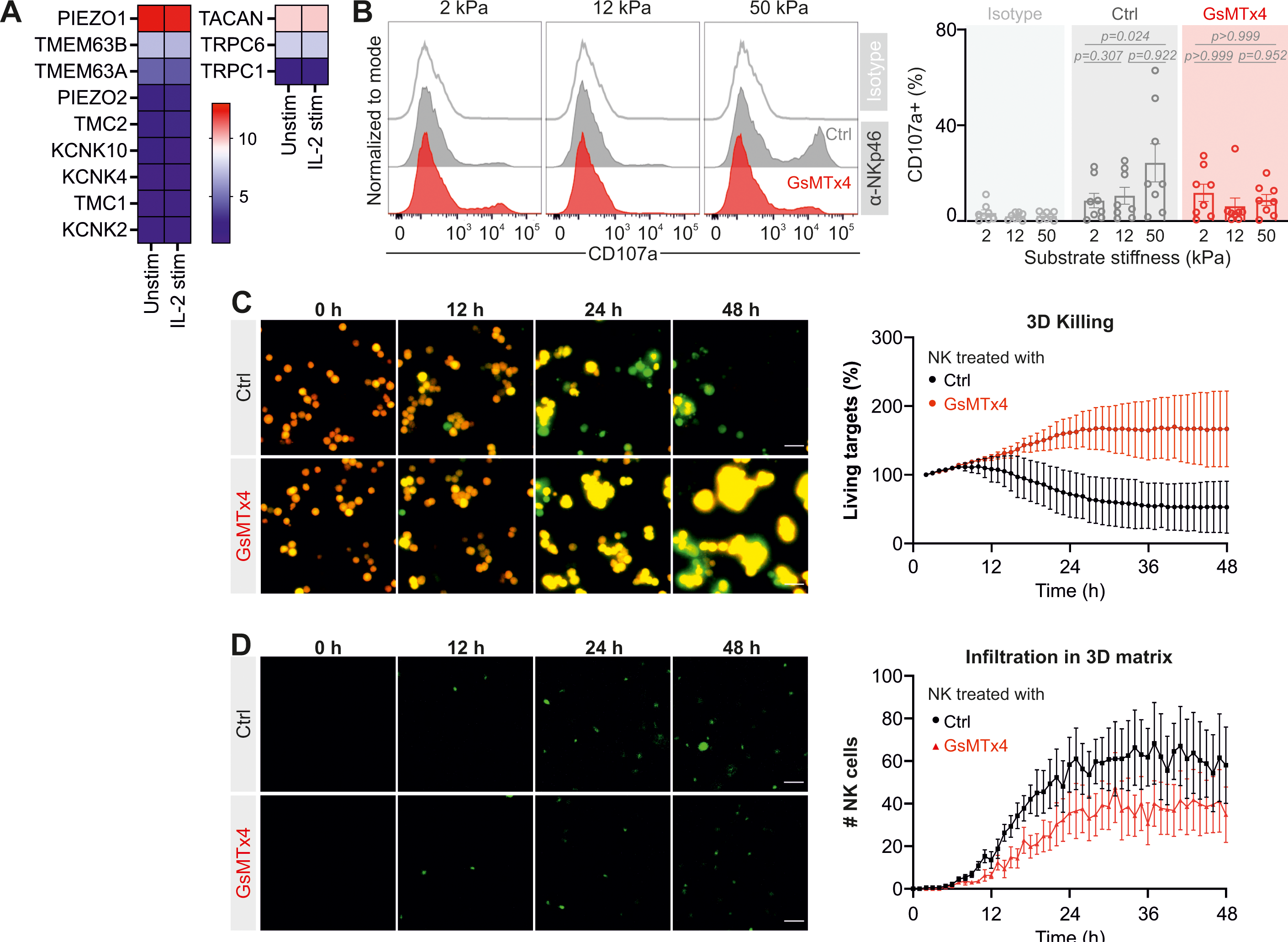
Inhibition of PIEZO1 reduces NK cell killing efficiency in 3D. Primary human NK cells from healthy donors were stimulated with IL-2 for three days. (**A**) Heatmap for the expression of mechanosensitive ion channels in unstimulated and IL-2 stimulated NK cells. The analysis is based on previously published microarray data (45). (**B**) GsMTx4 (50 µM) treatment abolishes substrate stiffness-dependent NK cell responsiveness. The activation of NK cells was evaluated using the CD107a degranulation assay. One representative donor is shown in the left panel and the quantification of all donors (n=8) is shown in the right panel. The Friedman test with Dunn’s multiple comparisons test was used for statistical analysis. (**C**) The killing efficiency of GsMTx4-treated NK cells in 3D is impaired. K562-pCasper target cells were embedded in collagen matrices (2 mg/ml) and the NK cells were added from the top. Live target cells are in orange-yellow color and apoptotic target cells in green. Time lapse of one representative donor is shown in the left panel and the quantification of all donors (n=3) is shown in the right panel. (**D**) GsMTx4-treated NK cells exhibit enhanced capability of infiltration into collagen matrices. The NK cells were stained with CFSE and added on the top of solidified collagen matrices (2 mg/ml). The NK cells approached the bottom were visualized (left panel) and quantified (right panel, n=4). A 20x magnification was used to obtain the images. The data are presented as mean ± SEM. Scale bars are 40 µm.

To investigate the effects of PIEZO1 perturbation on NK cell killing function, we further evaluated NK cell killing efficiency in a 3D scenario. Results from the 3D killing assay showed that GsMTx4-treated NK cells exhibited a substantially reduced killing efficiency compared to the control group (Fig. 3C, Supplementary Movie 3). Stiffness of the target cells was not affected by the presence of GsMTx4 (Supplementary Fig. 4A). To explore the underlying mechanisms, we examined the lytic granule pathway and NK cell migration. We found that GsMTx4 treatment did not alter the expression of cytotoxic proteins such as perforin and granzyme B (Supplementary Fig. 4B). Furthermore, degranulation induced by target cell recognition was even slightly enhanced (Supplementary Fig. 4C). Notably, the numbers of NK cells that infiltrated the 3D collagen matrix were greatly reduced after GsMTx4 treatment (Fig. 3D, Supplementary Movie 4). Aside from PIEZO1, GsMTx4 also targets a few other mechanically activated ion channels such as TRPC1, TRPC6 and TACAN. Although the expression levels of TRPC1, TRPC6 and TACAN are very low, if not negligible, compared to PIEZO1 (Fig. 3A). These observations suggest that PIEZO1-mediated mechanosensing is crucial for NK cells to execute their killing function in 3D mainly by regulating of NK infiltration into the 3D matrix.

To further validate the effect of PIEZO1, we utilized a PIEZO1 specific agonist, Yoda-1 (24) (Supplementary Fig. 5). Indeed, Yoda-1-treatment of NK cells accelerated killing kinetics in 3D collagen matrices with a significant reduction in the initiation time of killing events (Fig. 4A-B, Supplementary Movie 5). No difference was observed in degranulation induced by target cell recognition (Supplementary Fig. 6A), or in the conjugation between NK cells and target cells (Supplementary Fig. 6B) for Yoda-1-treated NK cells. Interestingly, infiltration of NK cells into 3D collagen matrix was substantially accelerated by Yoda-1 treatment (Fig. 4C-D, Supplementary Movie 6). Vehicle-treated NK cells first appeared in the focal plane at 6.6±1.6 hours, whereas first Yoda-1-treated NK cells approached the focal plane at around 4.8±1.7 hours (Fig. 4D). In addition, in a matrix-free environment, Yoda-1 treatment enhanced NK killing efficiency against both non-treated and softened tumor cells (Supplementary Figure 7A, B), while inhibition of PIEZO1 using GsMTx4 hindered NK killing against stiffened target cells (Supplementary Figure 7C), indicating that PIEZO1 can directly regulate the killing capacity of NK cells. Our results collectively suggest that PIEZO1 regulates both the NK killing processes and infiltration capability into 3D matrix, thereby fine-tuning the ultimate outcomes of tumor cell elimination.

**Figure 4.**
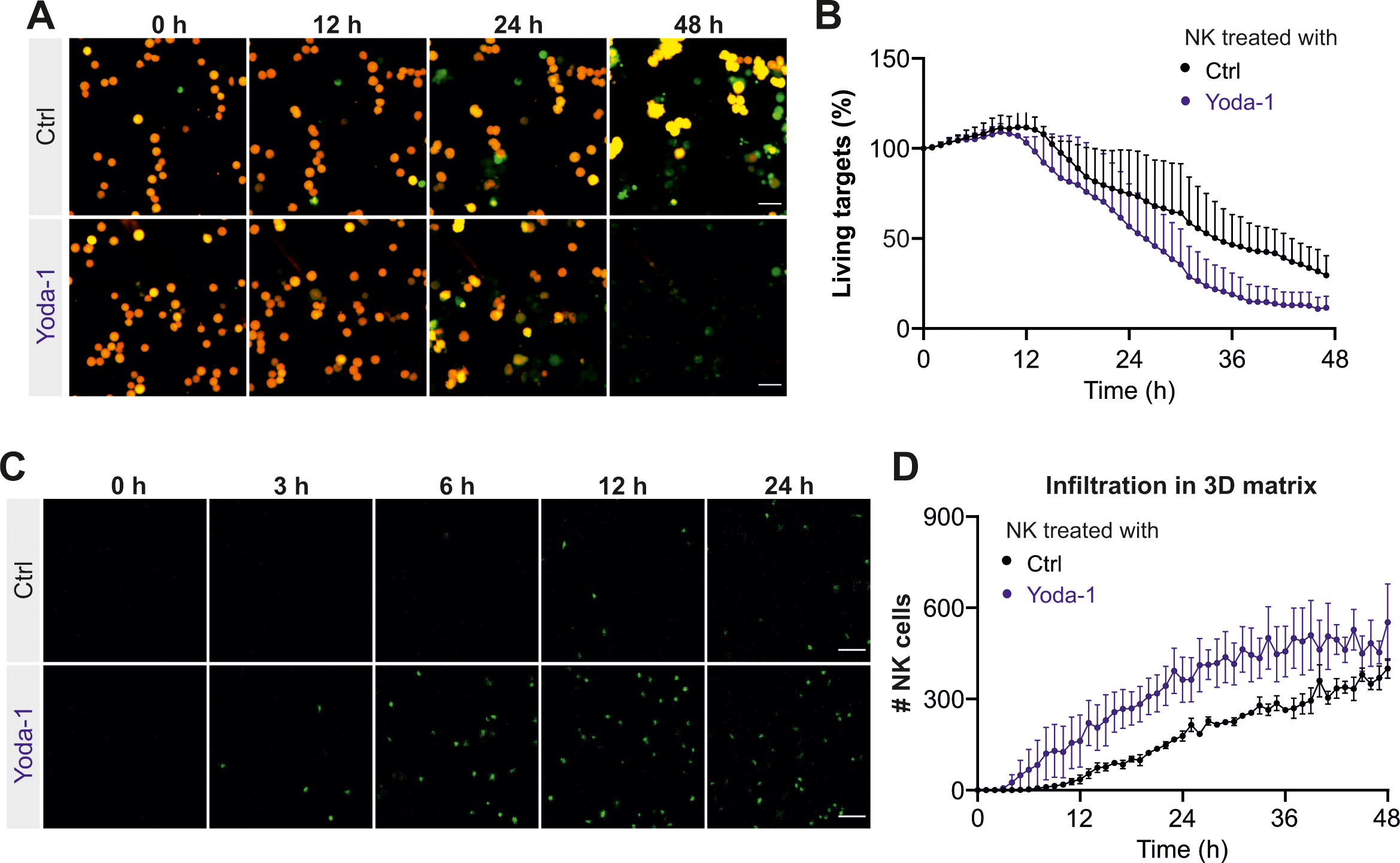
PIEZO1 activation enhances NK cell killing efficiency in 3D. Primary human NK cells from healthy donors were stimulated with IL-2 for three days. Yoda-1 (1 µM) was present in the medium during the experiments. (**A-B**) Yoda-1 treatment enhances the killing efficiency of NK cells in 3D. K562-pCasper target cells were embedded in collagen matrices (2 mg/ml) and the NK cells were added from the top. Live target cells are in orange-yellow color, and apoptotic target cells in green. Time lapse of one representative donor is shown in **A** and the quantification of all donors (n=3) is shown in **B**. (**C-D**) NK infiltration into 3D collagen matrices is enhanced by Yoda-1 treatment. The NK cells were stained with CFSE and added on the top of solidified collagen matrices (2 mg/ml). The NK cells approached the bottom were visualized (**C**) and quantified (**D**, n=3). A 20x magnification was used to obtain the images. The data are presented as mean ± SEM. Scale bars are 40 µm.

## Discussion

In our study, we have demonstrated that the killing efficiency of NK cells can be modulated by manipulating the stiffness of target cells. Cell softening is a recently discovered characteristic of malignant tumor cells, which is associated with tumorigenicity and malignancy. A rich body of evidence proves that cancer cells are softer than their non-malignant normal counterparts (25). For example, Cancerous breast epithelial cells are more deformable than their normal counterparts as determined by optical stretching (26). Ovarian cancer cells have a Young’s modulus in the range of 0.5-1 kPa, whereas their non-malignant counterpart have a stiffness of around 2 kPa, as determined by atomic force microscopy (9). Similarly, cervical cancer cells have an elastic modulus of ∼2 kPa, which is lower than that of normal human cervix epithelial cells (elastic modulus E ∼4-5 kPa) (10). Notably, even among malignant cells, the stiffness can vary, and softer cancer cells exhibit enhanced tumorigenicity, metastasis, and stemness. For example, among two ovarian cancer cell lines from the same specimen, the soft cells (HEY A8, ∼0.5 kPa) are more invasive than their stiffer counterparts (HEY, ∼0.9 kPa) (9). Soft cancer cells (breast cancer and melanoma cells, ∼0.2-0.3 kPa) require only ten cells to generate metastatic tumors in the lungs, whereas even 100 stiff cancer cells (∼0.8-1 kPa) are unable to produce any detectable lung metastasis (12). Softer cancer cells not only form more colonies with bigger sizes *in vitro*, but also have a substantially higher frequency of forming tumors *in vivo* (12). Stemness-associated genes are also up-regulated in soft tumor cells (12). Additionally, a study using cancer organoids embedded in 3D collagen has shown that cancer cells at the peripheral region are softer than the cells in the core region, and softer cancer cells are more invasive and metastatic (13). Tumor cells can be further softened by migration through confined spaces (13). In our study, the stiffness of non-treated tumor cells (∼1 kPa), softened tumor cells (∼0.6-0.7 kPa), or stiffened tumor cells (∼1.2-1.4 kPa) falls within the range of physiological stiffness as reported in the aforementioned studies. Our results demonstrate that softening (or stiffening) tumor cells substantially reduces (or enhances) elimination by NK cells, providing direct evidence supporting the hypothesis that cell softening is a mechanism by which malignant cells evade immune surveillance.

How does softening or stiffening of tumor cells affect the cytotoxicity of NK cells? In the process of cell killing, several key steps are critical, such as the initiation of IS formation, lytic granule enrichment and release, cytotoxic protein uptake by target cells, and detachment of NK cells after killing. Our study shows that softening or stiffening of tumor cells does not significantly alter lytic granule release, indicating that the events upstream of lytic granule release, such as IS formation and lytic granule enrichment, are unlikely to be significantly affected. However, we observed that the duration required to induce apoptosis or necrosis of tumor cells is prolonged for softened tumor cells compared to their stiff counterpart. Perforin-mediated pore formation on the plasma membrane of target cells is a critical step for directly lysing target cells or facilitating granzyme entry into target cells to induce apoptosis. Reduced tension of target cells impairs perforin-mediated pore formation and perforin-dependent killing (27). Therefore, to kill softened tumor cells, NK cells likely need to release more perforin or require more time to form the pores. Both scenarios require a longer duration for the killing process.

Following a successful killing, NK cells must detach from the dying or dead target cells in a timely manner to search for other targets and carry out more killing. Intriguingly, our observations show that the duration from the initiation of target cell apoptosis to the detachment of NK cells from stiffened target cells is substantially shorter than that from their softer counterparts. This finding suggests that alteration in tumor cell stiffness may influence the process of NK cell detachment. It is reported that conjugation with newly identified target cells can accelerate NK cell detachment from old target cells (28) and failed NK cell killing is linked to extended contact times (29, 30). Thus, the shortened contact times we observed for stiffened tumor cells possibly owe to more efficient NK cell killing. Studies on cytotoxic T lymphocytes suggest that recovery of cortical actin at the IS is essential to terminate lytic granule secretion, suggestively enabling or promoting T cells to detach from their target cells (31). PKCθ is required to break the symmetry of the IS, allowing naive T cells to disengage from their target cells (32). Additionally, calcium influx in T cells and apoptotic contraction also contribute to T cell disengagement from a target cell (33) (34). Therefore, it is possible that NK cells employ similar mechanisms to terminate killing processes and disassemble the IS, which is necessary for detachment from target cells. Interestingly, cell stiffness changes or increases after cell death (35, 36), which can serve as a direct cue to initiate NK cell detachment.

Recent studies have revealed that mechanical cues, particularly stiffness, can regulate the functions of NK cells. When primary human NK cells are stimulated with IL-2 on MICA-functionalized substrate with varying stiffness (30, 150, and 3000 kPa), they exhibit a bell-shaped response, with the maximum degranulation and clustering of DAP10 (an adaptor molecule downstream of NKG2D) occurring at 150 kPa (37). The application of mechanical forces to NK cells via MICA-functionalized nanowires (diameter ∼ 50 nm) enhances lytic granule degranulation upon NKG2D activation (38). Similarly, stiffer sodium alginate beads (34 and 254 kPa) functionalized with NKp30 antibody can induce full NK cell activation characterized by MTOC translocation and lytic granule polarization, whereas softer beads (9 kPa) failed to do so (7). Our data also suggest that NK cells exhibit greater degranulation triggered by activating receptors on stiffer hydrogels (12 and 50 kPa) compared to soft hydrogels (2 kPa) for most donors. However, for some donors, the levels of degranulation were comparable across all three stiffness levels. This variability is not associated with different expression levels of PIEZO1, as PIEZO1 expression levels are in a comparable range for various donors. Our findings suggest that NK cell responsiveness to stiffness is donor-dependent and may be attributed to variations in the expression of additional effector molecules involved in mechanosensing or transduction.

Mechanical cues are detected by various professional mechanosensors, primarily mechano-sensitive ion channel families, among which are the PIEZO family, TREK/TRAAK K2P (two-pore potassium) channels, TMEM63 (hyperosmolality-gated calcium-permeable) channels, and TMC (Transmembrane channel-like) 1/2 (14, 39). In our study, we report that PIEZO1 is the predominant mechanosensitive channel expressed in NK cells, indicating its indispensable role in mechanotransduction in NK cells. We observed that inhibiting PIEZO1 using GsMTx4 nearly abolished NK cell responsiveness to different substrate stiffness, greatly impaired NK cell-mediated cytotoxicity, and substantially reduced NK cell infiltration into 3D collagen matrices. Conversely, activating PIEZO1 with Yoda-1 enhanced the killing efficiency and infiltration capacity of NK cells. These findings demonstrate that PIEZO1-mediated mechanosensing is crucial for NK killing functions, highlighting PIEZO1 as a promising target to modulate NK functions, particularly in the context of solid tumors.

## Materials and Methods

### Antibodies and reagents

The following antibodies were purchased from BioLegend: PerCP anti-human CD3, BV421 anti-human CD3, APC anti-human CD56, BV421 anti-human CD107a, Biotin anti-human NKp46 (CD335), Biotin Mouse IgG1-κ Isotype, BV421 anti-human perforin, PE anti-human perforin, and PE anti-human granzyme B. Calcein-AM and carboxyfluorescein succinimidyl ester (CFSE) were purchased from Thermo Fischer Scientific, GsMTx4 from Smartox Biotechnology, blebbistatin from Cayman Chemical, Yoda-1 from Tocris, and Fibricol Collagen solution (10mg/ml Bovine Type I) from Advanced Biomatrix.

### Cell culture

Human peripheral blood mononuclear cells (PBMCs) of healthy donors were isolated from Leucocyte Reduction System Chamber using a gradient centrifugation method with Lymphocyte Separation Medium 1077 (PromoCell). Primary NK cells were isolated from the PBMCs using human NK cell isolation kit (Miltenyi) and then cultured in AIM V media with 10% FCS in presence of recombinant human IL-2 (100 U/ml, Miltenyi) for 3 days, unless mentioned otherwise. The purity is higher than 96%. As for K562 cells and K562-pCasper cells, they were cultured in RPMI medium supplemented with 10% FCS and 1 % penicillin and streptomycin (Thermo Fischer Scientific). For K562-pCasper cells, which stably express a FRET based apoptosis reporter (21), the culture medium was additionally supplemented with puromycin (0.2 µg/ml) (VWR).

### Preparation and biofunctionalization of hydrogels

Poly (acrylamide-co-acrylic acid) (PAAm-co-AA) hydrogels of varying stiffness were prepared and functionalized as previously described (19). Briefly, AAm monomer and bis-AAm crosslinker were mixed in different ratios, maintaining a constant ratio of AA. Hydrogel discs were prepared between two coverslips. PAAm-co-AA hydrogels were firstly functionalized with biotin-PEG8-NH2 by an EDC/NHS activation step as follows. The PAAm-co-AA film was covered with 100 µL EDC/NHS solution (39 mg/12 mg in 0.1 M, pH 4.5 MES buffer) for 15 min, washed thoroughly with PBS and directly incubated with 100 µL of biotin-PEG8-NH2 (1 mg/mL) solution in a petri dish for 2 h at RT. The functionalized hydrogels were washed with PBS 3 times and kept in PBS at 4°C until use. Functionalized hydrogels of varying stiffness were then incubated with streptavidin solution (100 µL, 100 μg/mL) for 1 –1.5 h (2 kPa) or 2.5 –3 h (12 and 50 kPa) according to a previously reported protocol (19). Streptavidin-functionalized hydrogels were then incubated with biotinylated anti-NKp46 (100 μg/mL, 30 μL) or IgG isotype (100 μg/mL, 30 μL) overnight at 4°C.

### CD107a degranulation assay

To assess stimulation-induced NK degranulation, NK cells were either settled on substrates functionalized with NKp46 antibody or incubated with K562 cells in presence of BV421 anti-CD107a antibody and protein transport inhibitor Golgi stop (BD Biosciences) at 37°C with 5% CO_2_ for 4 hours. Then the cell suspension was stained with PerCP anti-human CD3, APC anti-human CD56 antibodies at 4°C in dark for 30 minutes. The samples were analyzed using FACSVerse (BD Biosciences). The CD3^-^CD56^+^ population was gated for NK cells. FlowJo v10 (FLOWOJO, LLC) was used for data analysis.

### Determination of NK cell killing kinetics

To determine NK killing efficiency in 3D environments, the assay was conducted as described previously (40). Briefly, K562-pCasper target cells were resuspend in neutralized bovine collagen I (2 mg/ml) and plated in a black 96-well plate with flat clear bottom (Corning/Merck) at a density of 25,000 cells/40 µl per well. The plate was centrifuged to spin down the target cells on the bottom and then polymerized at 37°C with 5 % CO_2_ for 1 hour. NK cells were added on the top of the collagen matrix with an E:T ratio of 1:1 if not otherwise specified. A high content imaging system ImageXpress (Molecular Devices) was used to acquire images at 37°C with 5 % CO_2_ every 20 minutes for 48 hours. K562-pCasper target cells with a FRET signal above the threshold (maximal FRET signals in GFP-positive target cells) were taken as live target cells. The number of live target cells at each time point was normalized to that at time 0. AIMV medium supplemented with 10% FCS was used in this assay. The images were processed and analyzed using ImageJ.

To determine NK killing capacity in a matrix free 2D scenario, K562-pCasper target cells were settled in a 96-well half area microplate with clear flat bottom (Corning/Merck) at a density of 25,000 cells per well at RT for 20 min. Subsequently, NK cells were added from above with an E:T ratio of 0.5:1. Killing events were visualized using ImageXpress at 37°C with 5 % CO_2_ every 20 minutes for 8 hours.

### NK cell migration in 3D collagen matrices

NK cells were stained with CFSE (5 µM in PBS/4.5% FCS) at room temperature for 15 minutes, washed once with PBS, then resuspended in AIMV/10% FCS and kept at 37°C with 5 % CO_2_ overnight for recovery. Sample preparation for light-sheet microscopy was describe previously (41). Briefly, CFSE-stained NK cells were resuspended in neutralized bovine collagen I (2 mg/ml), and this cell suspension was polymerized in a capillary at 37°C with 5 % CO_2_ for 1 hour. Subsequently, the sample was mounted in the sample chamber filled with RPMI medium. Z-stacks (step size ∼ 2 µm for ∼ 100 slices) were acquired using a Z.1 light-sheet microscope (Zeiss) at 37°C every 30 seconds for 30 minutes. Imaris 8.1.2 (Bitplane) was used to automatically track fluorescently labeled NK cells to quantify cell velocity and persistence.

### Analysis of NK cell infiltration into 3D collagen matrices

NK cells were stained with CFSE (5 µM in PBS/4.5% FCS) at room temperature for 15 minutes, washed once with PBS, then resuspended in AIMV/10% FCS and kept at 37°C with 5 % CO_2_ overnight for recovery. Neutralized bovine collagen I (2 mg/ml) was plated 40 µl per well in a black 96-well plate with flat clear bottom (Corning/Merck). The plate was kept at 37°C with 5 % CO_2_ for 1 hour. After solidification, CFSE-stained NK cells (25,000 cells/well) were added on the top of the matrix. To identify the position of the bottom, one well plated with NK cells without collagen was used as reference. Images focused on the bottom were acquired using ImageXpress at 37°C with 5 % CO_2_ every 20 minutes for 48 hours. ImageJ was used to identify fluorescently labeled NK cells and quantify the number of infiltrated NK cells for each time point.

### Determination of cytotoxic protein expression

NK cells were washed twice with PBS containing 0.5% BSA, then stained with PerCP anti-human CD3, APC anti-human CD56 antibodies at 4°C in dark for 30 minutes. To assess the expression of perforin and granzyme B, these pre-stained NK cells were fixed with pre-chilled 4% paraformaldehyde (PFA) for 20 minutes, permeabilized with PBS containing 0.1% saponin, 0.5% BSA, and 5% FCS at room temperature for 10 minutes, then stained with BV421 anti-human Perforin and PE anti-human Granzyme B antibodies at room temperature in dark for 40 minutes. The samples were analyzed using FACSVerse (BD Biosciences). The CD3^-^CD56^+^ population was gated for NK cells. FlowJo v10 (FLOWOJO, LLC) was used for data analysis.

### Real-time deformability cytometry (RT-DC)

To assess the stiffness of K562pCasper cells RT-DC (Zellmechanik Dresden) was used (42). K562pCasper were either treated with DMSO or Blebbistatin for 12 hours after which they were resuspended in 100 µl of Cell Carrier B solution (phosphate-buffered saline with the addition of long-chain methylcellulose polymers of 0.6 w/v%). A microfluidic PDMS chip with a 300 µm long central constriction of 30 µm×30 µm cross-section was assembled on the stage of an inverted microscope (Zeiss). The cell suspension was loaded on the chip using a syringe pump. The cells flowing through the microfluidic channel deform due to the shear stresses and pressure gradient caused by the flow profile (43). Each event is imaged live using a CMOS camera and analysed in real-time. At least 3000 events were acquired per condition at a flowrate of 0.16 µls^-1^. The mechanical properties of the cells were analyzed using ShapeOut 2 (Zellmechanik Dresden) which employs linear mixed models to calculate statistical significances.

### Immunostaining

NK cells stimulated with 100 U/ml IL-2 for 3 days were seeded on the Poly-L-ornithine coated coverslip for 15 min at RT, then were fixed with 4% paraformaldehyde and permeabilized with 0.1% Triton-100 in PBS at RT for 15 min, followed by a blocking step with 2% BSA in PBS at RT for 1 hr. Samples were incubated with anti-PIEZO1 antibody at 4°C overnight and then with Alex488 goat anti-rabbit secondary antibody. F-actin and nucleus are labeled with phalloidin and Hoechst 33342, respectively. The images were acquired by Cell Observer wide-field microscopy with 40× oil objective (1.3 NA) with a step size of 0.3 µm for z-stacks. The acquired images were deconvolved using the Huygens Essential Software.

### Ca^2+^ imaging

Ca^2+^ imaging was carried out as described previously (44). Briefly, Jurkat T cells were loaded with Fura-2-AM (1 μM) at RT for 25 min, resuspended in 1 mM Ca^2+^ Ringer’s solution, then seeded on a poly-L-ornithine coated coverslip. Live-cell imaging was acquired excitation of 340 nm, 380 nm and infrared every 5 sec at RT. After the first 30 cycles, measurements were paused and Yoda (1µM in 1 mM Ca^2+^ Ringer’s solution) was perfused into the chamber, then the measurements were resumed. This time point was defined as Time 0. The images were analyzed using T.I.L.L. Vision software.

### Ethical considerations

Research carried out for this study with material from healthy donors (Leukocyte Reduction System Chambers from human blood donors) has been authorized by the local ethic committee of the “Ärztekammer des Saarlandes” (Identification Nr. 84/15, Prof. Dr. Rettig-Stürmer).

### Statistical analysis

GraphPad Prism 8.3Software (GraphPad) was used for statistical analysis. For RT-DC, linear mixed models were used. For other statistical analyses, D’Agostino and Pearson test was used to test the normality. Between two paired groups, paired t-test was used for normal distribution and Wilcoxon matched-pairs signed rank test was used for non-normal distribution. Between two unpaired groups, unpaired t-test was used for normal distribution and Mann-Whitney-U-test was used for non-normal distribution. To compare three or more groups, one-way ANOVA test was used, and multiple comparisons were done with Dunn’s multiple comparisons test.

## Author Contributions

AKY performed most of the experiments and the corresponding analysis if not mentioned otherwise; JZ prepared hydrogels; XZ performed real-time killing assay and the analysis, and helped with image processing, figures and movies; GM, FL, DB, and OO helped with RT-DC; RZ performed immunostaining and Ca^2+^ imaging; SS examined purity of NK cells and PIEZO1 expression; MH and AdC helped with data interpretation and provided critical feedback on all aspects of the project; BQ generated concepts and designed experiments; All authors contributed to the writing, editing and cross-checking of the manuscript.

## Data Availability Statement

The data that support the findings of this study are available from the corresponding author upon reasonable request.

## Acknowledgments

We thank the Institute for Clinical Hemostaseology and Transfusion Medicine for providing donor blood, Carmen Hässig, Cora Hoxha, Gertrud Schäfer, Kathrin Förderer for excellent technical help, Lea Kaschek for assisting compositing images, Mohamed Hamed and Eva C. Schwarz for assisting analysing microarray data. This project was funded by the Deutsche Forschungsgemeinschaft (SFB 1027 Project A2 to BQ, A10 to FL, A11 to MH, B6 to AdC; and Forschungsgroßgeräte (GZ: INST 256/423-1 FUGG for the flow cytometer to MH, GZ: INST 256/429-1 FUGB for ImageXpress to MH, and GZ: INST 256/569-1 for the RT-DC to FL and BQ).

## Conflict of Interest Statement

The authors declare that the research was conducted in the absence of any commercial or financial relationships that could be construed as a potential conflict of interest.

## Notes

### Competing Interest Statement

The authors have declared no competing interest.

### Summary of Updates

The title has been corrected to match the uploaded files

## References

1. S. Sivori et al., Human NK cells: surface receptors, inhibitory checkpoints, and translational applications. Cell Mol Immunol 16, 430–441 (2019).

2. J. S. Orange, Formation and function of the lytic NK-cell immunological synapse. Nat Rev Immunol 8, 713–725 (2008).

3. K. Krzewski, J. E. Coligan, Human NK cell lytic granules and regulation of their exocytosis. Front Immunol 3, 335 (2012).

4. M. W. Pickup, J. K. Mouw, V. M. Weaver, The extracellular matrix modulates the hallmarks of cancer. EMBO Rep 15, 1243–1253 (2014).

5. C. Rianna, M. Radmacher, S. Kumar, Direct evidence that tumor cells soften when navigating confined spaces. Mol Biol Cell 31, 1726–1734 (2020).

6. A. B. Roberts et al., Tumor cell nuclei soften during transendothelial migration. J Biomech 121, 110400 (2021).

7. D. Friedman et al., Natural killer cell immune synapse formation and cytotoxicity are controlled by tension of the target interface. J Cell Sci 134 (2021).

8. O. Matalon et al., Actin retrograde flow controls natural killer cell response by regulating the conformation state of SHP-1. EMBO J 37 (2018).

9. W. Xu et al., Cell stiffness is a biomarker of the metastatic potential of ovarian cancer cells. PLoS One 7, e46609 (2012).

10. K. Hayashi, M. Iwata, Stiffness of cancer cells measured with an AFM indentation method. J Mech Behav Biomed Mater 49, 105–111 (2015).

11. Y. L. Han et al., Cell swelling, softening and invasion in a three-dimensional breast cancer model. Nat Phys 16, 101–108 (2020).

12. J. Lv et al., Cell softness regulates tumorigenicity and stemness of cancer cells. EMBO J 40, e106123 (2021).

13. C. Rianna, M. Radmacher, S. Kumar, Direct evidence that tumor cells soften when navigating confined spaces. Molecular Biology of the Cell 31, 1726–1734 (2020).

14. J. M. Kefauver, A. B. Ward, A. Patapoutian, Discoveries in structure and physiology of mechanically activated ion channels. Nature 587, 567–576 (2020).

15. J. M. Hope et al., Fluid shear stress enhances T cell activation through Piezo1. BMC Biol 20, 61 (2022).

16. C. S. C. Liu et al., Cutting Edge: Piezo1 Mechanosensors Optimize Human T Cell Activation. J Immunol 200, 1255–1260 (2018).

17. A. Jairaman et al., Piezo1 channels restrain regulatory T cells but are dispensable for effector CD4(+) T cell responses. Sci Adv 7 (2021).

18. A. G. Solis et al., Mechanosensation of cyclical force by PIEZO1 is essential for innate immunity. Nature 573, 69–74 (2019).

19. J. Zhang et al., Micropatterned soft hydrogels to study the interplay of receptors and forces in T cell activation. Acta Biomater 119, 234–246 (2021).

20. G. Alter, J. M. Malenfant, M. Altfeld, CD107a as a functional marker for the identification of natural killer cell activity. J Immunol Methods 294, 15–22 (2004).

21. C. S. Backes et al., Natural killer cells induce distinct modes of cancer cell death: Discrimination, quantification, and modulation of apoptosis, necrosis, and mixed forms. J Biol Chem 293, 16348–16363 (2018).

22. C. J. Chan et al., Myosin II Activity Softens Cells in Suspension. Biophys J 108, 1856–1869 (2015).

23. C. Bae, F. Sachs, P. A. Gottlieb, The mechanosensitive ion channel Piezo1 is inhibited by the peptide GsMTx4. Biochemistry 50, 6295–6300 (2011).

24. R. Syeda et al., Chemical activation of the mechanotransduction channel Piezo1. Elife 4 (2015).

25. C. Alibert, B. Goud, J. B. Manneville, Are cancer cells really softer than normal cells? Biol Cell 109, 167–189 (2017).

26. J. Guck et al., Optical deformability as an inherent cell marker for testing malignant transformation and metastatic competence. Biophys J 88, 3689–3698 (2005).

27. R. Basu et al., Cytotoxic T Cells Use Mechanical Force to Potentiate Target Cell Killing. Cell 165, 100–110 (2016).

28. P. Netter, M. Anft, C. Watzl, Termination of the Activating NK Cell Immunological Synapse Is an Active and Regulated Process. J Immunol 199, 2528–2535 (2017).

29. M. R. Jenkins et al., Failed CTL/NK cell killing and cytokine hypersecretion are directly linked through prolonged synapse time. J Exp Med 212, 307–317 (2015).

30. M. Anft et al., NK cell detachment from target cells is regulated by successful cytotoxicity and influences cytokine production. Cell Mol Immunol 17, 347–355 (2020).

31. A. T. Ritter et al., Cortical actin recovery at the immunological synapse leads to termination of lytic granule secretion in cytotoxic T lymphocytes. Proc Natl Acad Sci U S A 114, E6585–E6594 (2017).

32. T. N. Sims et al., Opposing effects of PKCtheta and WASp on symmetry breaking and relocation of the immunological synapse. Cell 129, 773–785 (2007).

33. A. Bohineust, Z. Garcia, H. Beuneu, F. Lemaitre, P. Bousso, Termination of T cell priming relies on a phase of unresponsiveness promoting disengagement from APCs and T cell division. J Exp Med 215, 1481–1492 (2018).

34. E. E. Sanchez et al., Apoptotic contraction drives target cell release by cytotoxic T cells. Nat Immunol 24, 1434–1442 (2023).

35. N. I. Nikolaev, T. Muller, D. J. Williams, Y. Liu, Changes in the stiffness of human mesenchymal stem cells with the progress of cell death as measured by atomic force microscopy. J Biomech 47, 625–630 (2014).

36. M. Islam et al., Microfluidic Sorting of Cells by Viability Based on Differences in Cell Stiffness. Sci Rep 7, 1997 (2017).

37. L. Mordechay et al., Mechanical Regulation of the Cytotoxic Activity of Natural Killer Cells. ACS Biomater Sci Eng 7, 122–132 (2021).

38. G. Le Saux et al., Nanoscale Mechanosensing of Natural Killer Cells is Revealed by Antigen-Functionalized Nanowires. Adv Mater 31, e1805954 (2019).

39. D. Douguet, E. Honore, Mammalian Mechanoelectrical Transduction: Structure and Function of Force-Gated Ion Channels. Cell 179, 340–354 (2019).

40. R. Zhao, A. K. Yanamandra, B. Qu, A high-throughput 3D kinetic killing assay. 10.1101/2023.04.11.536280 (2023).

41. R. Schoppmeyer, R. Zhao, M. Hoth, B. Qu, Light-sheet Microscopy for Three-dimensional Visualization of Human Immune Cells. J Vis Exp 10.3791/57651 (2018).

42. O. Otto et al., Real-time deformability cytometry: on-the-fly cell mechanical phenotyping. Nat Methods 12, 199–202, 194 p following 202 (2015).

43. A. Mietke et al., Extracting Cell Stiffness from Real-Time Deformability Cytometry: Theory and Experiment. Biophys J 109, 2023–2036 (2015).

44. B. Qu et al., Docking of lytic granules at the immunological synapse in human CTL requires Vti1b-dependent pairing with CD3 endosomes. J Immunol 186, 6894–6904 (2011).

45. S. Zophel et al., Identification of molecular candidates which regulate calcium-dependent CD8(+) T-cell cytotoxicity. Mol Immunol 157, 202–213 (2023).

