## Supplementary Information for "PIEZO1-mediated mechanosensing governs NK cell killing efficiency and infiltration in three-dimensional matrices"

### Supplementary Figures

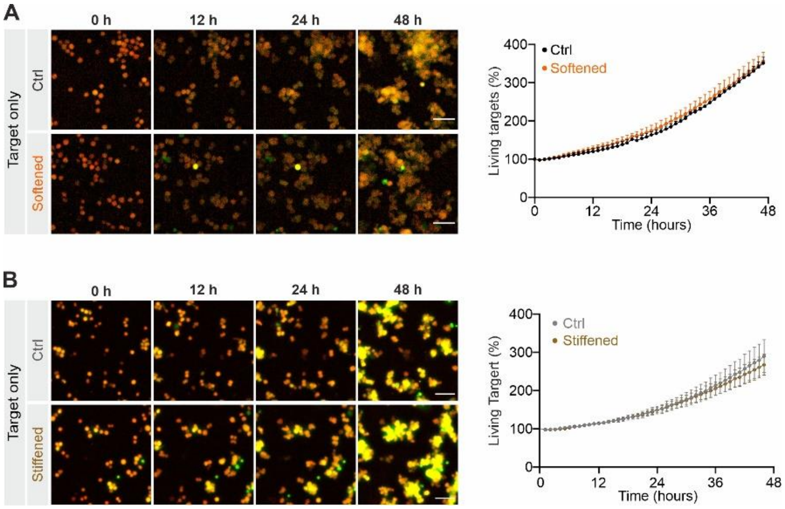


**Supplementary Figure 1. Proliferation of target cells is not affected by changes in stiffness.** Primary human NK cells from healthy donors were stimulated with IL-2 for three days prior to the experiments. K562-pCasper target cells were pre-treated with either DMSO (Softened) or blebbstatin (Stiffened) and then embedded in collagen matrices (2 mg/ml). Live target cells are in orange-yellow color and apoptotic target cells in green. Time lapse of killing events obtained in 20x magnification. The data are presented as mean ± SEM. Scale bars are 40 µm.

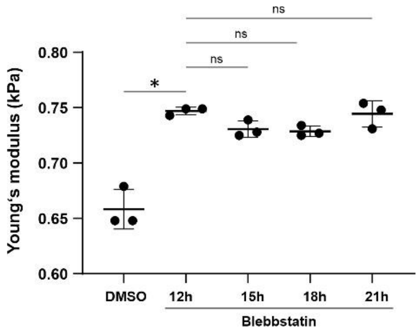


**Supplementary Figure 2. Stiffness of blebbstatin-treated target cells.** K562-pCasper cells were treated with blebbstatin. The corresponding stiffness was determined using RT-DC at the indicated time points. Results are shown as mean±SEM, from three independent experiments. The Kruskal-Wallis test was used for statistical analysis.

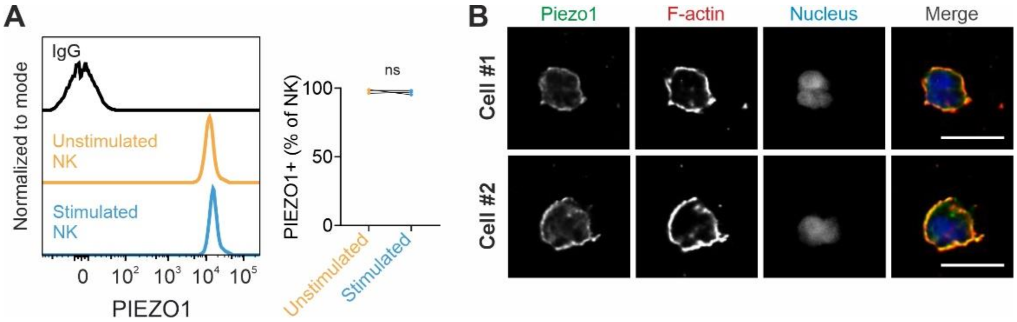


**Supplementary Figure 3. PIEZO Expression at the protein level.** (**A**) PIEZO1 expression levels were examined using flow cytometry for unstimulated and IL-2-stimulated primary human NK cells. Results are from three donors. The Student's paired t-test was used for statistical analysis. (**B**) Distribution of PIEZO1 in NK cells. Primary human NK cells from healthy donors were stimulated with IL-2 for three days prior to the experiments. PIEZO1, F-actin, and the nucleus was indicated with anti-PIEZO1 antibody, phalloidin, and Hoechst 33342, respectively. Two exemplary cells out of two independent experiments are shown. Scale bars are 10 µm.

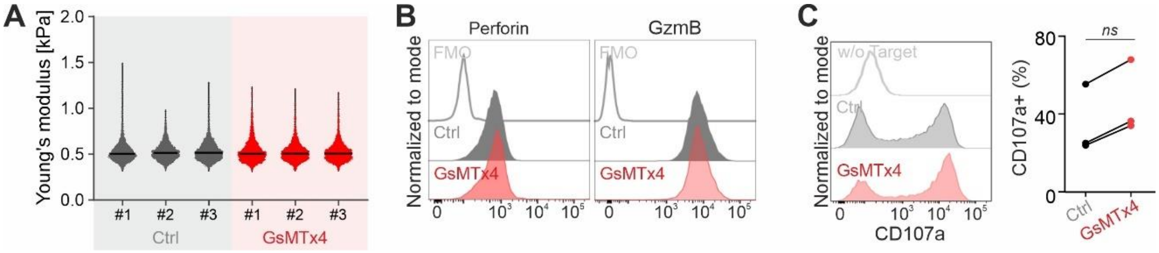


**Supplementary Figure 4. Effect of GsMTx4 on target cells and NK cells.** (**A**) The presence of GsMTx4 does not affect target cell stiffness. K562-pCasper target cells were treated with GsMTx4 (50 µM) at 37°C with 5% $CO_2$ overnight. Cell stiffness was determined using RT-DC. The results of three independent experiments are shown. (**B-C**) GsMTx4 treatment does not affect lytic granule pathway in NK cells. Primary human NK cells from healthy donors were stimulated with IL-2 for three days. The expression of perforin and granzyme B was assessed using flow cytometry (**B**, n=2). NK cell degranulation induced by target cell recognition was evaluated using the CD107a degranulation assay (**C**, n=3).

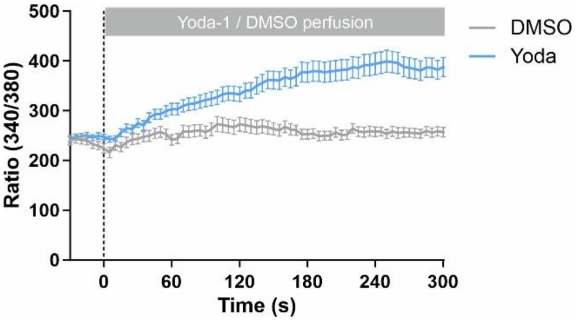


**Supplementary Figure 5. Yoda-1 activates PIEZO1.** Jurkat T cells were loaded with Fura-2 and seeded on a poly-L-ornithine coated coverslip for $Ca^{2+}$ imaging. Live-cell imaging was carried out at RT every 5 sec for three channels: 340 nm, 380 nm and infrared. Results are shown as mean±SEM, from three independent experiments.

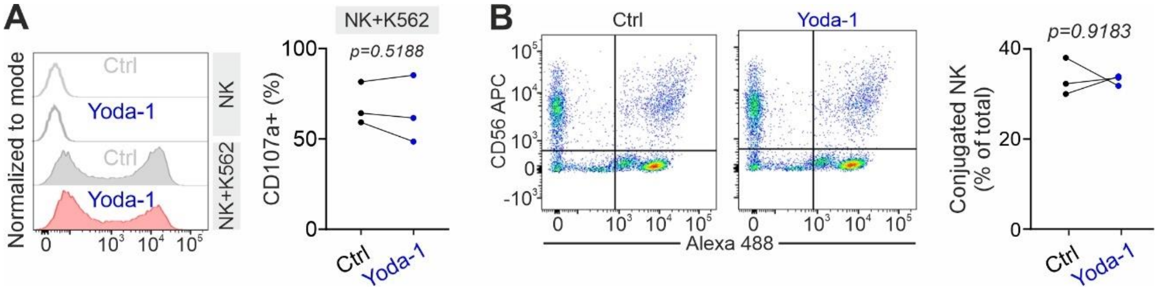


**Supplementary Figure 6. Yoda-1 treatment does not affect degranulation and NK/target conjugation.** NK cell degranulation induced by target cell recognition was evaluated using the CD107a degranulation assay (**A**, n=3). To evaluate NK/target conjugation, the NK cells stained with anti-CD56 and K562 stained with Calcein-AM were incubated at 37°C for 10 minutes. Double positive doublets were taken as NK/target conjugates and the analysis is shown in **B**.

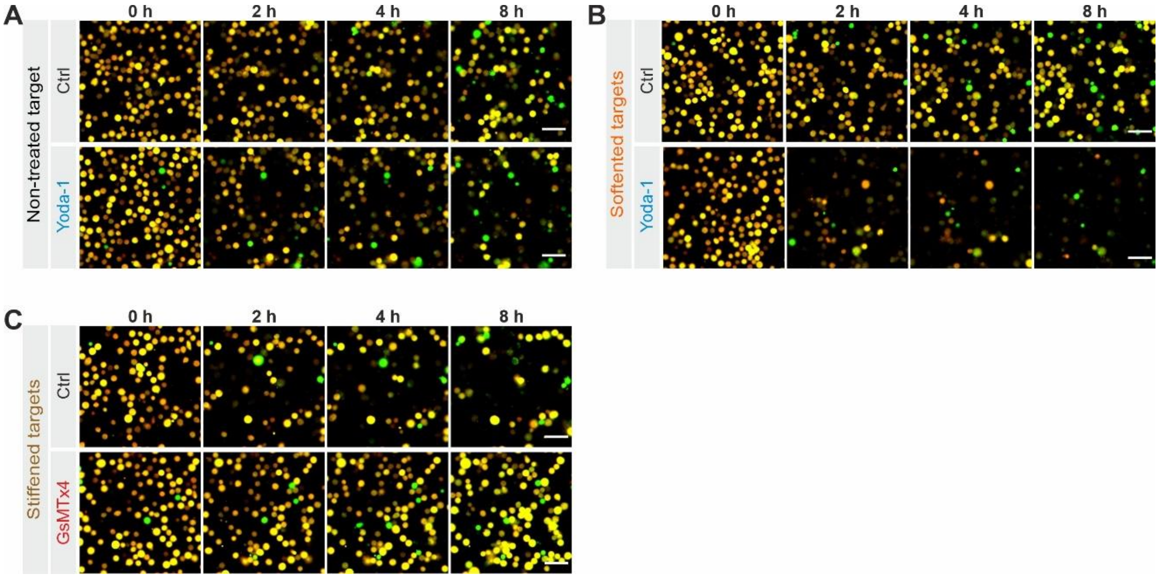


**Supplementary Figure 7. PIEZO1 activity regulates NK cell killing capacity.** Primary human NK cells from healthy donors and K562-pCasper target cells were co-cultured in a flat-bottomed 96-well plate for live-cell imaging. Target cells were non-treated (A), softened with DMSO (B), or stiffened with blebbstatin (C). Yoda-1 (1 µM) or GsMTx4 (50 µM) was present in the medium during the experiments. Live target cells are in orange-yellow color, and apoptotic target cells in green. Time lapse of one representative donor out of three is shown. A 20x magnification was used to obtain the images. Scale bars are 40 µm.

### Movie legends

**Movie 1. Softening tumor cells impairs NK cell killing efficiency in 3D.** Primary human NK cells from healthy donors were stimulated with IL-2 for three days. K562-pCasper cells were pre-treated with DMSO (1:2000, Softened) for 12 hours. Untreated K562-pCasper cells were used as the Ctrl group. Time lapse of one representative donor is shown (n=3). Live target cells are in orange-yellow color and apoptotic target cells in green. Scale bars are 40 µm.

**Movie 2. NK cells eliminate stiffened tumor cells more efficiently.** Primary human NK cells from healthy donors were stimulated with IL-2 for three days. K562-pCasper cells were pre-treated with Blebbistatin (50 µM, Stiffened) or vehicle control for 12 hours. Live target cells are in orange-yellow color and apoptotic target cells in green. Time lapse of one representative donor is shown (n=3). Scale bars are 40 µm.

**Movie 3. The killing efficiency of GsMTx4-treated NK cells in 3D is impaired.** Primary human NK cells from healthy donors were stimulated with IL-2 for three days. K562-pCasper target cells were embedded in collagen matrices (2 mg/ml) and the NK cells were added from the top. Live target cells are in orange-yellow color and apoptotic target cells in green. Time lapse of one representative donor is shown (n=3). Scale bars are 40 µm.

**Movie 4. GsMTx4-treated NK cells exhibit enhanced capability of infiltration into collagen matrices.** Primary human NK cells from healthy donors were stimulated with IL-2 for three days. The NK cells were stained with CFSE and added on the top of solidified collagen matrices (2 mg/ml). The NK cells approached the bottom were visualized. One representative donor is shown (n=4). Scale bars are 40 µm.

**Movie 5. Yoda-1 treatment enhances the killing efficiency of NK cells in 3D.** Primary human NK cells from healthy donors were stimulated with IL-2 for three days. K562-pCasper target cells were embedded in collagen matrices (2 mg/ml) and the NK cells were added from the top. Live target cells are in orange-yellow color, and apoptotic target cells in green. Time lapse of one representative donor is shown (n=3). Scale bars are 40 µm.

**Movie 6. NK infiltration into 3D collagen matrices is enhanced by Yoda-1 treatment.** Primary human NK cells from healthy donors were stimulated with IL-2 for three days. The NK cells were stained with CFSE and added on the top of solidified collagen matrices (2 mg/ml). The NK cells approached the bottom were visualized. Time lapse of one representative donor is shown (n=3). Scale bars are 40 µm.
